# Serine Starvation Silences Estrogen Receptor Signaling through Histone Hypoacetylation

**DOI:** 10.1101/2021.09.05.459037

**Authors:** Albert M. Li, Yang Li, Bo He, Haowen Jiang, Yaniel Ramirez, Meng-Ning Zhou, Chao Lu, Joshua J. Gruber, Erinn B. Rankin, Jiangbin Ye

## Abstract

Estrogen receptor (ER) plays important roles in regulating normal development and female reproductive system function. Loss of ER pathway activity is a hallmark of breast cancer progression, associated with accelerated tumor proliferation and resistance to endocrine therapy. How ER loss occurs remains poorly understood. Here, we show that serine starvation, a metabolic stress often found in solid tumors, downregulates estrogen receptor alpha (ERα) expression, represses transcriptional targets such as progesterone receptor (PR), and reduces sensitivity to antiestrogens, suggesting a transition of ER-positive (ER^+^) breast cancer cells to an ER/PR-negative (ER^-^/PR^-^) state. ER downregulation under serine starvation is accompanied by a global loss of histone acetylation. These chromatin changes are driven by metabolic reprogramming triggered by serine starvation, particularly lower glucose flux through glycolysis and the TCA cycle, leading to reduced acetyl-CoA levels and histone hypoacetylation. Supplementation with acetate or glycerol triacetate (GTA), precursors of acetyl-CoA, restores H3K27 acetylation and ERα expression under serine starvation. Therefore, a major consequence of serine starvation in breast cancer could be global chromatin changes that influence lineage-specific gene expression.

## INTRODUCTION

Breast cancer is the leading cause of death in women. The etiology of this heterogeneous disease remains poorly understood, with numerous studies devoted to elucidating the genetic and cell-of-origin drivers of this disease. The loss of estrogen receptor (ER) signaling represents a crucial step in the progression to aggressive breast cancer, as patients who present with ER^+^ tumors typically respond well to endocrine therapies such as tamoxifen or fulvestrant. Some refractory tumors lose ER expression^1^; this could be attributed to either transcriptional repression of ER or the outgrowth of a small population of ER^-^ cells present in the initial tumor^2^. Consistent with the latter model, a different cell of origin in the mammary epithelium has been postulated to give rise to each different breast cancer subtype. ER^+^ (Luminal A/B) tumors are thought to be derived from mature luminal cells, whereas ER^-^ (claudin-low and basal-like) tumors are thought to be derived from basal or luminal progenitor-like cells^3^. More recent data has challenged this model, suggesting that luminal progenitors could be the common cell of origin for both luminal (ER^+^) and basal-like (ER^-^) breast cancers^4^. As such, it is still unclear if and how the tumor microenvironment (TME) plays a role in the switch between ER^+^ and ER^-^ cancer cell states, and how this switch is transcriptionally regulated. Understanding how TME factors such as metabolic stress contribute to ER loss could shape current strategies for therapeutic breast cancer interventions.

Altered metabolism is a hallmark of cancer cells^5^. Recent work has elucidated some of the metabolic vulnerabilities of breast cancer subtypes^6-9^, but how metabolic reprogramming occurs is still poorly understood. The amino acid serine is depleted in the core of melanoma tumors^10^, pancreatic tumors^11^, and in the mammary fat pad^12^, the tissue of origin for breast cancer. Because serine is essential for the synthesis of macromolecules such as proteins, nucleotides, and lipids^13^, serine starvation slows cancer cell proliferation^14,15^. At the molecular level, serine starvation diverts glucose carbons to *de novo* serine synthesis while upregulating the key serine synthesis enzymes phosphoglycerate dehydrogenase (PHGDH), phosphoserine amino transferase (PSAT1), and phosphoserine phosphatase (PSPH)^16^. Intriguingly, PHGDH is upregulated in about 70% of ER^-^ breast cancers compared to ER^+^ tumors^17^, and luminal breast cancer cell lines tend to be more auxotrophic for serine compared to basal-like breast cancer cell lines^18^. These observations raise the intriguing possibility that a common driver may underlie the simultaneous gain of serine synthesis pathway and ER silencing in breast cancer.

The present work identifies serine availability as a mediator of ER status in breast cancer cells. Serine starvation elevates the expression of enzymes involved in serine synthesis and the mitochondrial serine and one-carbon (1C) unit pathway, represses ER signaling and transcriptional activity, and promotes resistance to endocrine therapy. Acetate supplementation restores chromatin state and rescues ERα mRNA and protein expression, highlighting metabolic interventions as a potential strategy to prevent acquired resistance and manipulate lineage-associated cell states in breast cancer.

## RESULTS

### Serine starvation upregulates serine biosynthesis and mitochondrial serine catabolism genes while repressing estrogen receptor pathway signaling

Our previous study identified upregulation of mitochondrial serine and 1C unit pathway enzymes in metastatic subclones derived from the triple-negative breast cancer (TNBC) cell line MDA-MB-231^7^. Many of the enzymes involved in the mitochondrial catabolism of serine are transcriptionally induced by serine starvation, so we hypothesized that serine-limiting conditions, which may occur during passage in immunodeficient mice^19-21^, may serve as a trigger to reprogram gene expression in MDA-MB-231 cells towards a metastatic gene expression signature. Indeed, in parental 231 cells, acute serine starvation (24h treatment) transcriptionally elevated enzymes involved in the *de novo* serine synthesis pathway (PHGDH, PSAT1, and PSPH), as well as those in the mitochondrial serine and 1C unit pathway (SHMT2, MTHFD2, MTHFD1L) (Figure S1A). In contrast, this induction was absent in 4175-LM (lung metastatic subclone) cells that already expressed higher levels of these enzymes. Consistent with higher enzyme expression, we observed higher *de novo* serine synthesis from glucose in 1833-BoM (bone metastatic subclone) cells under serine starvation, as evidenced by higher labeling of M+3 serine (Figure S1B). These results suggest that serine starvation could be a driver of higher serine catabolism and synthesis gene expression in more aggressive breast cancer cells.

To uncover other pathways affected by serine starvation that may lead to more aggressive cancer cell phenotypes, we performed global transcriptome profiling via RNA sequencing (RNA-seq) in parental MDA-MB-231 cells after 24h of serine starvation. Pathway analysis of differentially expressed genes using the ENRICHR gene list enrichment tool revealed numerous upregulated and downregulated pathways, including the late estrogen response pathway (Figure 1A and Figure S1C). This was surprising because the MDA-MB-231 cell line is a triple-negative breast cancer cell line and is thought to lack estrogen receptor protein expression. We next performed a similar analysis in the estrogen receptor positive (ER^+^) cell line MCF7. Serine starvation induced a robust transcriptional upregulation of genes involved in serine synthesis and the mitochondrial serine and 1C unit pathway (Figure S1D). In this cell line, we also observed a significant scoring of the late estrogen response pathway among top downregulated pathways (Figure 1B). Comparative Gene Set Enrichment Analysis (GSEA) of expression data taken from both the ER^-^ MDA-MB-231 and the ER^+^ MCF7 cell lines cultured in complete vs. serine-free media also revealed an enrichment of genes in the late estrogen response pathway (Figure 1C, 1D), including estrogen receptor alpha (ERα, encoded by *ESR1*) and its transcriptional targets *KRT19* and *PGR* (Figure 1E). The silencing of *ESR1* and *PGR* upon serine starvation was validated using q-PCR (Figure 1F). To determine whether ESR1 repression was associated with endocrine resistance, we treated MCF7 cells and another ER^+^ cell line, T47D, with fulvestrant (ICI) and observed reduced sensitivity of serine-starved cells to the anti-proliferative effects of fulvestrant (Figure 1G). Reduced anti-proliferative effects were also observed when cells were treated with a related estrogen antagonist, tamoxifen (4OHT), under serine starvation (Figure S2A). Together, these data suggest that serine starvation reprograms cell to an ER^-^ like, estrogen-independent state.

**Figure 1.**
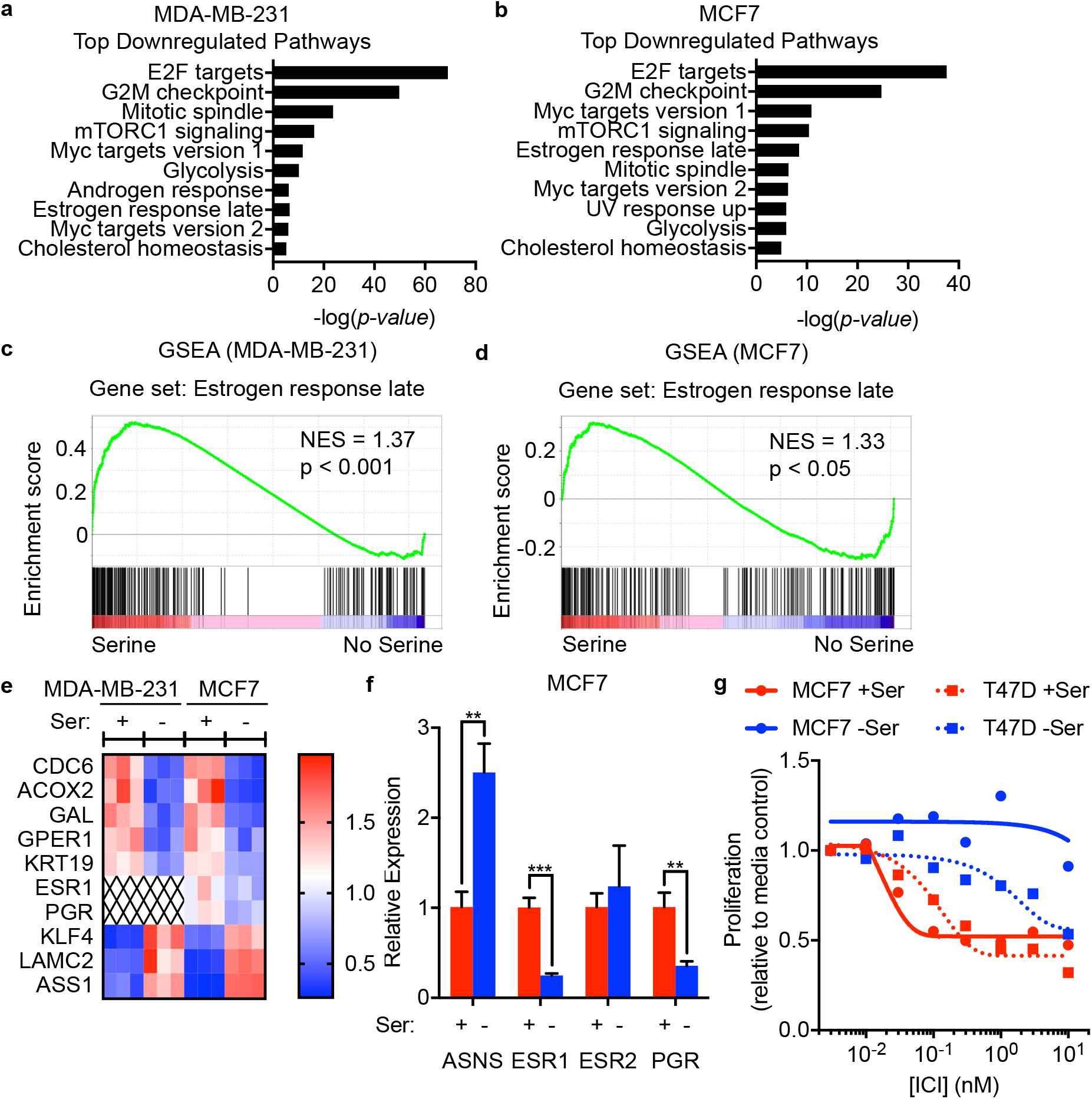
Serine starvation represses the estrogen signaling pathway and promotes endocrine resistance. **(a-d)** Pathway analysis of top downregulated pathways via ENRICHR **(a**,**b)** and enrichment plots via GSEA **(c**,**d)** on RNA-seq performed in control vs. 24h serine-starved MDA-MB-231 **(a**,**c)** or MCF7 **(b**,**d)** cells. -log(p-value) represents the degree of pathway significance. NES, nuclear enrichment score. **(e)** Heatmap of the expression of select estrogen signaling pathway related genes as measured by RNA-seq in (a-d). **(f)** Quantitative RT-PCR analysis for estrogen signaling pathway genes in 24h serine-starved MCF7 cells. (**g**) Four day proliferation assay on MCF7 and T47D cells in control (+Ser) or serine-free (-Ser) media, treated with increasing doses of fulvestrant (ICI). Data represent mean ± SD **(f, g)**, N = 3 per group. Statistics: two-tailed unpaired Student’s t test; **p < 0.01, ***p < 0.001.

### Rapid global histone hypoacetylation under serine starvation drives ER*α* loss

Given that chromatin state regulates gene transcription, we next tested how chromatin markers changed in response to serine starvation. In MCF7 cells, withdrawal of serine from the cell culture media resulted in a time-dependent decreases in nuclear ERα expression, with the most significant repression at 24h starvation (Figure 2A). This pattern was most closely mirrored by a reduction in global H3K27ac levels. Analysis of nuclear extracts from T47D cells also revealed a time-dependent loss of ERα and H3K27ac levels (Figure S2B). Interestingly, the repressive H3K9me3 and H3K27me3 marks were elevated under serine starvation (Figure 2A). To determine whether histone hypoacetylation could be a cause of ERα silencing, we treated serine-starved cells with the HDAC inhibitors suberanilohydroxamic acid (SAHA) and romidepsin (Rom). Both SAHA and Rom treatment elevated H3K27ac levels in MCF7 cells (Figure 2B). Importantly, at the mRNA level, we observed the complete restoration of *ESR1* and its transcriptional target *PGR* under serine starvation (Figure 2C). We thus reasoned that the mechanism underlying estrogen receptor silencing involves loss of H3K27ac.

**Figure 2.**
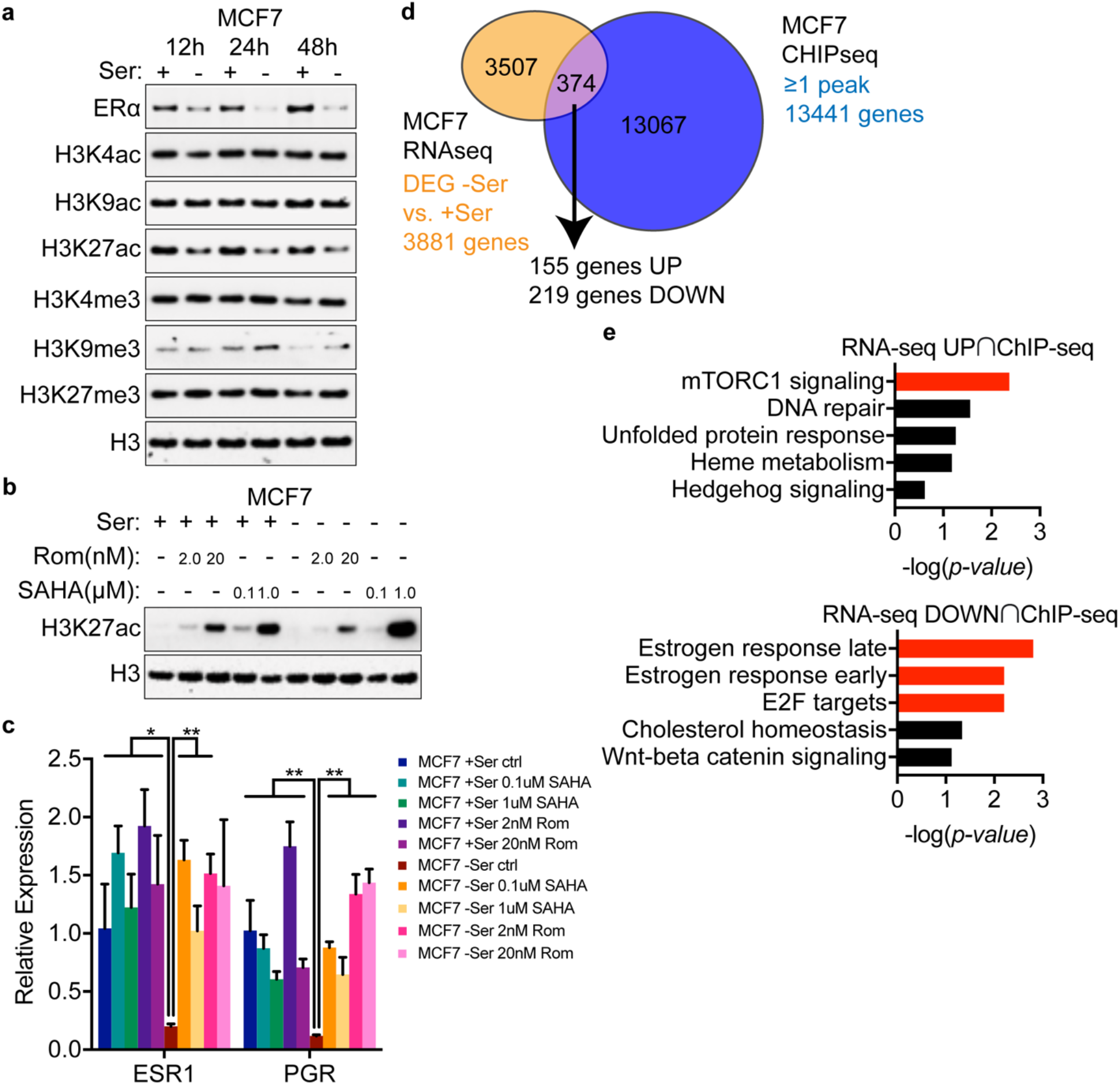
Serine starvation suppresses estrogen receptor signaling through H3K27 hypoacetylation. **(a)** Immunoblot against estrogen receptor alpha (ERα) and common acetylation and methylation H3K residues, performed on nuclear fraction lysates isolated from MCF7 cells grown in +/- Ser media at 12h, 24h, and 48h. Band intensities were quantified with Imagelab 6.0.1 software (Bio-Rad) and normalized to the loading control (H3) and the first sample (+Ser 12h) of each blot. **(b, c)** Immunoblot for H3K27ac in nuclear lysates **(b)** and quantitative RT-PCR for *ESR1* and *PGR* **(c)** from MCF7 cells grown in +/- Ser media treated with either suberanilohydroxamic acid (SAHA) or romidepsin (Rom) for 18h. **(d)** Venn diagram of overlap between differentially expressed genes (DEGs) in +/-Ser cultured MCF7 cells (from RNA-seq) and genes containing at least one H3K27ac peak (from ChIP-seq). **(e)** Pathway analysis of top upregulated or downregulated pathways via ENRICHR on overlapped genes identified in **(d)**. Significant pathways (p < 0.01) are represented by red bars. Data are mean ± SD **(c)**, N = 3 per group. Statistics: two-tailed unpaired Student’s t test; *p < 0.05, **p < 0.01.

To further explore the relationship between serine starvation-mediated transcriptional changes and global H3K27ac dynamics, we analyzed a previously published MCF7 H3K27ac dataset^22^ generated by chromatin immunoprecipitation followed by sequencing (ChIP-seq). We identified 13,441 genes containing at least one H3K27ac peak within or around gene bodies (Figure 2D, Figure S2C). We found 374 differentially enriched genes (DEGs) in serine-starved MCF7 cells also harbored the H3K27ac modification. Strikingly, ENRICHR pathway analysis on the subset of 219 downregulated genes revealed an enrichment in the Hallmark estrogen response late and early pathways that was absent in the same analysis performed on the subset of 155 upregulated genes (Figure 2E, Figure S2D). These results further support the idea that genes in the ER pathway are preferentially sensitive to serine starvation-mediated H3K27ac loss.

### Serine starvation reduced glucose flux to the TCA cycle and depletes intracellular citrate and Ac-CoA levels

We next sought to understand the metabolic changes that underlie loss of histone acetylation, as metabolism is known to provide the substrates, cofactors, and allosteric regulators for chromatin modifying enzymes^23-25^. We performed targeted metabolomics analysis of serine-starved cells at various time points, focusing on metabolites involved in glycolysis, shunt pathways, and the TCA cycle. Serine starvation induced a significant enrichment at early time points (1h and 2h) of upstream glycolytic intermediates, including fructose 1,6-bisphosphate (F16BP), glycerol 3-phosphate (G3P)/dihydroxyacetone phosphate (DHAP), and the serine precursor 3-phosphoglycerate (3PG) (Figure 3A, Figure S3A). Serine levels, however, rapidly reduced at 1h and failed to recover by 24h (Figure 3A, Figure S3A). We also observed lower levels of the glycolysis end products lactate and pyruvate, which feeds into the TCA cycle (Figure 3A, Figure S3A). Furthermore, the levels of the TCA cycle intermediates were reduced at early time points of serine starvation. Citrate and alpha ketoglutarate (αKG) levels returned to normal by 24h while succinate, fumarate, and malate levels remained low (Figure 3B, Figure S3B). Given that nuclear histone acetylation is dependent on cytosolic acetyl-CoA generation by ATP citrate lyase (ACLY)^26^, we hypothesize that reduced citrate levels under serine starvation could be the trigger for histone hypoacetylation.

**Figure 3.**
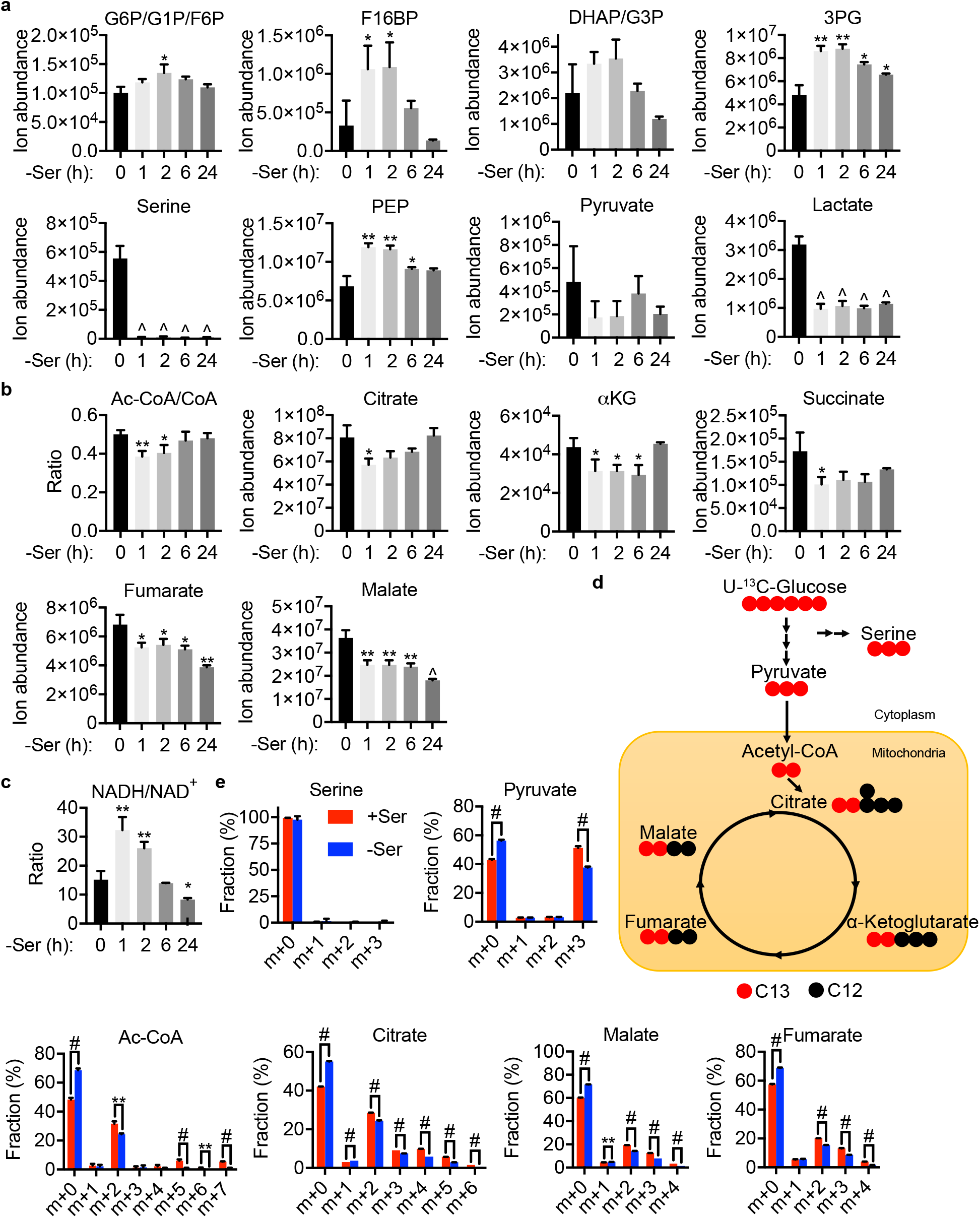
Reduced glucose flux through glycolysis and the TCA cycle under serine starvation leads to citrate and acetyl-CoA insufficiency. **(a-c)** LC-MS was used to determine the levels of intermediates in glycolysis **(a)**, the TCA cycle **(b)**, and the NADH/NAD^+^ ratio **(c)** at various time points of serine starvation in MCF7 cells. **(d)** Schematic of U-^13^C-glucose labeling to serine and intermediates in glycolysis and the TCA cycle. **(e)** Isotopomer distribution of serine, pyruvate, acetyl-CoA, citrate, fumarate, and malate from MCF7 cells cultured in control (+Ser) or serine-free (-Ser), glucose-free media supplemented with U-^13^C-glucose for 6h. Data are mean ± SD **(a-c, e)**, N = 3 per group. Statistics: two-tailed unpaired Student’s t test; *p < 0.05, **p < 0.01, ^p < 0.001, #p < 0.0001. Abbreviations: G6P, glucose 6-phosphate; G1P, glucose 1-phosphate; F6P, fructose 6-phoshpate; F16BP, fructose 1,6-bisphosphate; DHAP, dihydroxyacetone phosphate; G3P, glycerol 3-phosphate; 3PG, 3-phosphoglycerate; PEP, phosphoenolpyruvate; Ac-CoA, acetyl-Coenzyme A; CoA, coenzyme A; αKG, alpha ketoglutarate; NAD^+^, nicotinamide adenine dinucleotide.

Lower metabolite levels could be a consequence of increased consumption or reduced generation. To distinguish between the two possibilities, we performed isotope tracing with uniformly ^13^C-labeled glucose. Glucose-derived acetyl-CoA carries an isotope label mass heavy by 2 (M+2), which is passed on to subsequent intermediates in the TCA cycle upon incorporation with oxaloacetate into citrate (Figure 3D). Compared with cells grown in complete media, serine-starved MCF7 cells showed a reduction of M+2 labeled acetyl-coA and citrate, indicating reduced flux into the TCA cycle (Figure 3E). Directly upstream of carbon incorporation into the TCA cycle, M+3 labeled pyruvate levels were also reduced in serine-starved cells, suggesting lower glycolysis flux. The lower glucose flux through glycolysis and the TCA cycle was associated with a transient increase in the NADH/NAD^+^ ratio, which can affect rate-limiting steps of glycolysis such as the GAPDH reaction (Figure 3C, S3C). Seahorse assays further revealed multiple dysregulated parameters of mitochondrial respiration including lower basal respiration, in line with previous observations in serine/glycine-starved p53 wild-type colon cancer cells^14^ (Figure S3D, S3E). Consistent with the kinetics of rapid depletion of intracellular serine observed earlier (Figure 3A), M+3 serine was absent in MCF7 cells grown in either complete or serine-starved conditions (Figure 3E), suggesting a defect in *de novo* serine synthesis, possibly because the reductive stress (increased NADH/NAD^+^ ratio, Figure 3C) inhibited PHGDH activity^27^. Taken together, these results suggest a model whereby serine starvation reduces glucose flux to glycolysis and the TCA cycle, leading to reduced acetyl-CoA levels and histone hypoacetylation.

### Acetate rescues serine-starvation mediated histone hypoacetylation to maintain ERα expression

It was previously reported that the loss of histone acetylation at gene promoters under LDHA inactivation^28^ or hypoxia^29,30^ could be enhanced with acetate supplementation, thus restoring gene expression. Therefore, we hypothesize that the reduction of acetyl-CoA and histone acetylation triggered by serine starvation could be overcome by acetate supplementation, through either ACSS2 or the ACSS1/3 to ACLY pathway (Figure 4A). Indeed, supplementation with two different forms of acetate, sodium acetate (NaAc) or glyceryl triacetate (GTA), restored nuclear H3K27ac levels under serine starvation (Figure 4B). Importantly, GTA rescued *ESR1* mRNA, ERα protein levels, and ER signaling activity as measured by expression of the ERα transcriptional target *PGR* (Figure 4C, 4D). Taken together, these results further validate that serine starvation silences estrogen receptor signaling through an epigenetic mechanism of acetyl-CoA depletion-mediated transcriptional repression.

**Figure 4.**
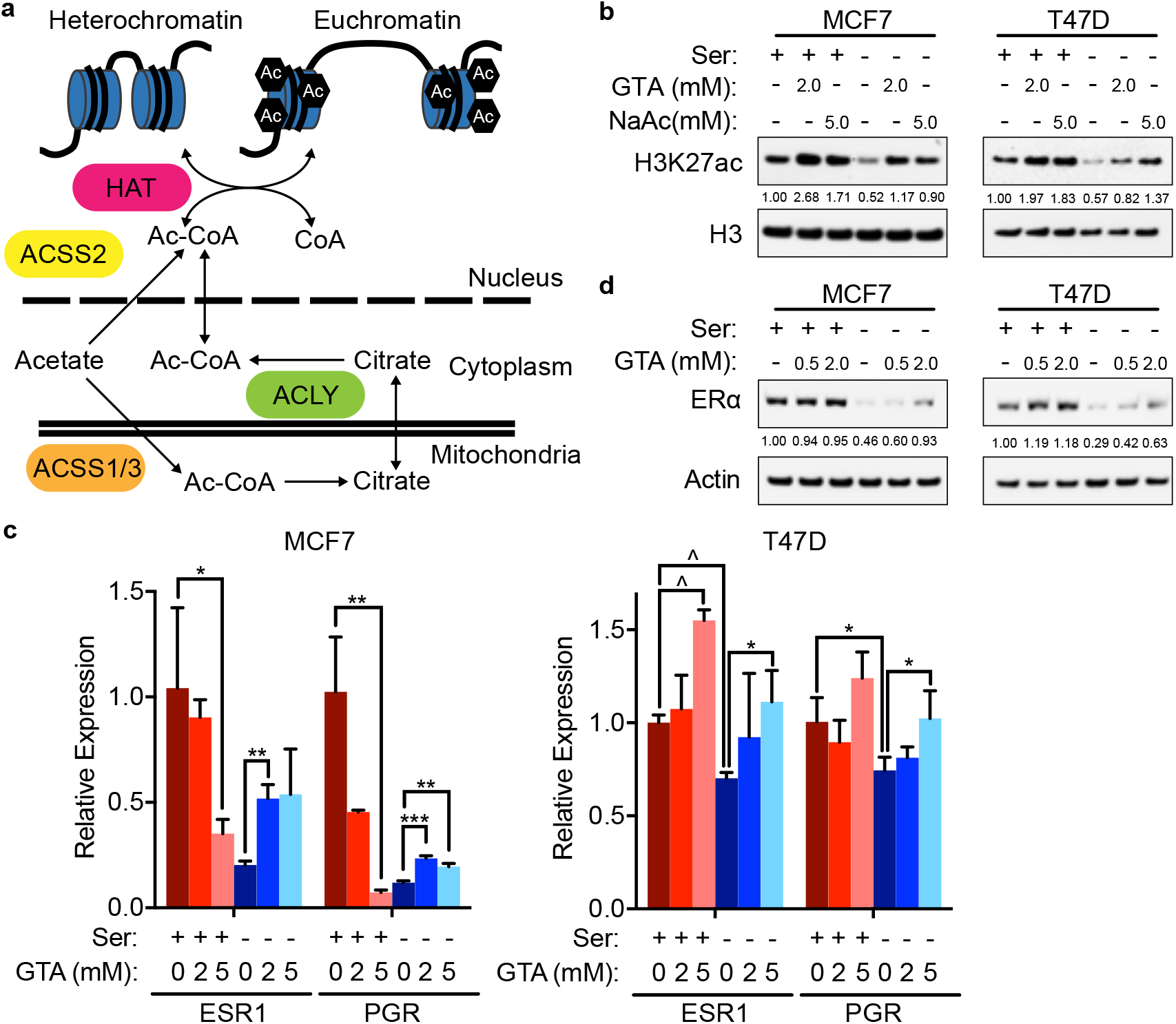
Acetate supplementation restores H3K27ac and estrogen receptor signaling. **(a)** Schematic of two ways acetate may contribute to nuclear Ac-CoA pools to support histone acetyltransferase (HAT) activity. **(b)** Immunoblot for H3K27ac in nuclear extracts from MCF7 and T47D cells grown in +/- Ser media supplemented with glycerol triacetate (GTA) or sodium acetate (NaAc) for 24h. (**c**) Quantitative RT-PCR analysis for *ESR1* and *PGR* in MCF7 and T47D cells grown in +/- Ser media supplemented with glycerol triacetate (GTA) for 18h. **(d)** Immunoblot for ERα in whole cell extracts from MCF7 and T47D cells grown in +/- Ser media supplemented with different concentrations of GTA for 24h. Data are mean ± SD **(c)**, N = 3 per group. Statistics: two-tailed unpaired Student’s t test; *p < 0.05, **p < 0.01, ***p < 0.001.

## DISCUSSION

Serine starvation has shown promising results in preclinical cancer models at improving survival^15^ and slowing primary tumor growth^12^. However, cancer cells adapt to low serine levels by rewiring glucose flux and transcriptionally activating enzymes involved in *de novo* serine synthesis^16^. Recent studies have linked higher *de novo* serine synthesis with cancer cell aggression and metastasis^31,32^. However, how low serine levels reprogram chromatin state and gene expression is unclear. In this study, we demonstrate that serine starvation reprograms epigenetics through metabolic rewiring. Serine-starved ER^+^ breast cancer cells lower glucose flux through glycolysis. This may be due to a transient increase in NADH/NAD^+^ ratios generated by the PHGDH reaction^27^, which could inhibit rate-limiting steps of glycolysis. The inability of ER^+^ cells to synthesize serine, as evidenced lack of M+3 serine from isotope-labeled glucose, further impedes glycolysis due to serine-dependent PKM2 activation^16,33^. While the net metabolic consequence of serine starvation is reduced lactate levels, masquerading as “Warburg effect” inhibition, the global epigenetic changes and the reduction of glucose flux to the TCA cycle mimic the effects of hypoxia^30^, which promotes the Warburg effect. Indeed, it was previously reported that hypoxia reduces both ESR1 mRNA and ERα protein levels^34^, akin to the reduction of ERα mRNA and protein levels in ER^+^ cells subjected to acute serine starvation.

Therefore, we believe that the key cell-autonomous consequence of the Warburg effect may not be lactate generation, but rather an inhibition of mitochondrial activity and TCA cycle activity, culminating in epigenetic reprogramming and the silencing of genes important for defining cell lineage. Our findings provide a potential explanation for the higher expression of serine synthesis genes in ER^-^ tumors compared to ER^+^ tumors—i.e., some serine synthesis^active^/ER^-^ tumors arose from serine-starvation-mediated ER silencing of serine synthesis^inactive^/ER^+^ tumors. Of note, while ER^-^ 1833-BoM subclone cells displayed robust *de novo* serine synthesis under serine starvation, both ER^-^ parental 231 cells and ER^+^ MCF7 cells failed to synthesize serine from glucose despite transcriptional upregulation of *PHGDH, PSAT1*, and *PSPH* (Figure S1A, S1B, S1D), suggesting that additional properties such as enhanced NAD^+^ generation^27,35^ may be necessary for breast cancer cells to fully acquire serine synthesis ability. Therefore, the overall reduction of glycolysis flux, rather than diversion of glucose carbons to serine, may be the main cause of serine starvation-mediated histone hypoacetylation and chromatin remodeling.

Can metabolic-epigenetic-driven silencing of ER be reversed? Our rescue experiments with acetate demonstrate that providing a metabolic substrate to restore histone acetylation may be key to preventing ER silencing under a breast tumor microenvironment stress. Given earlier observations that the combined inhibition of histone deacetylases and DNA methyltransferases with small-molecule inhibitors could restore ER expression in TNBC cells^36^, it is tempting to speculate the potential for metabolic interventions to prevent the transcriptional loss of ER during breast cancer progression. Future studies are required to further understand which metabolites are required to sensitize advanced ER^-^ tumors to endocrine therapy.

## Acknowledgements

This work was supported by a NIH T32 Training Grant (CA009302-40) to A.M.L., and an American Cancer Society Research Scholar Grant (RSG-20-036-01) and a Stanford Maternal and Child Health Research Institute Research Scholar Award (2020) to J.Y. We thank the members of the Rankin and Ye labs for general assistance and fruitful discussions.

## Author contributions

Conception and design: A.M.L., C.L., J.Y.

Development of methodology: A.M.L., Y.L., B.H., H.J., E.B.R., J.Y.

Acquisition of data: A.M.L., Y.L., B.H., H.J., Y.R., J.Y.

Analysis and interpretation of data: A.M.L., Y.L., B.H., H.J., M-N.Z., C.L., J.J.G., E.B.R., J.Y.

Writing, review, and/or revision of the manuscript: A.M.L., C.L., J.J.G., E.B.R., J.Y.

## STAR Methods

### Lead contact

Further information and requests for resources and reagents should be directed to and will be fulfilled by the lead contact, Jiangbin Ye.

### Materials availability

This study did not generate new unique reagents.

### Data and code availability

The serine starvation RNA-seq data will be deposited at GEO. This paper does not report original code. Any additional information required to reanalyze the data reported in this paper is available from the lead contact upon request.

## EXPERIMENTAL MODEL AND SUBJECT DETAILS

### Cell culture

MCF7, T47D, MDA-MB-231 parental, 4175-LM lung metastatic subclone, and 1833-BoM bone metastatic subclone cell lines were a kind gift from Dr. Joan Massagué (Memorial Sloan Kettering Cancer Center). All cell lines were cultured in DMEM/F12 (Caisson Labs) with 10% fetal bovine serum (Sigma) with 1% penicillin/streptomycin (Thermo Scientific). Cells were tested every 3 to 6 months and found negative for Mycoplasma (MycoAlert Mycoplasma Detection Kit; Lonza). All cell lines used were passaged no more than 10 times from the time of thawing. Cell lines were not authenticated.

## METHOD DETAILS

### Serine starvation and proliferation assay

Cell proliferation was assessed using a hemocytometer and light microscope. 5.0 × 10^4^ /well cells were plated in a 12-well plate and attached overnight. The media was changed the following day to glucose/serine/glycine-free media (Teknova) supplemented with 10% dialyzed fetal bovine serum (Sigma), 2.0 g/L glucose, and 0.13 mM glycine to reconstitute -Ser media, and with 0.3 mM serine to reconstitute +Ser media. The media from each well was collected prior to addition of Trypsin (Thermo Scientific) to adherent cells. Each well of trypsinized cells was combined with its corresponding collected media, spun down, then resuspended in 10% fetal bovine serum media prior to the addition of 0.4% Trypan Blue Stain (Thermo Scientific) at a volume of 1:1 to distinguish live from dead cells. Stock solutions of fulvestrant (ICI, Cayman) and tamoxifen (4OHT, MP Biomedicals) were reconstituted in DMSO and stored in aliquots at -20 °C prior to addition to drug treatment assays.

### Protein isolation, SDS-PAGE, and immunoblot

For whole cell lysates, cells were washed with ice cold PBS buffer and lysed with RIPA buffer (Thermo Scientific) supplemented with 1:100 Halt Protease and Phosphatase Inhibitor Cocktail (Thermo Scientific) for 15 minutes over ice. For nuclear fractionation, cells were washed with ice cold PBS buffer and lysed with harvest lysis buffer (10 mM HEPES pH 7.9, 50 mM NaCl, 500 mM sucrose, 0.1 mM EDTA, 0.5% Triton x 100) supplemented with 1:100 Halt Protease and Phosphatase Inhibitor Cocktail (Thermo Scientific) for 10 minutes over ice. Cell lysates were centrifuged at 5000 RPM for 5 minutes at 4 °C. Supernatants were transferred to a new Eppendorf tube and labeled as the “Cytosolic Fraction.” Insoluble pellets containing nuclear proteins, labeled as the “Nuclear Fraction,” were further lysed with nuclear lysis buffer (10 mM HEPES pH 7.9, 500 mM NaCl, 0.1 mM EDTA, 0.1 mM EGTA, 0.1% NP40) supplemented with 1:100 Halt Protease and Phosphatase Inhibitor Cocktail (Thermo Scientific) for 10 minutes over ice. Nuclear Fractions were further sonicated in a Biorupter Plus sonication device (Diagenode) at 4 °C for 15 cycles of 60 seconds ON, 30 seconds OFF on the High setting. Nuclear Fractions were then centrifuged at 15,000 RPM for 10 minutes at 4 °C. Protein concentrations were determined by the Pierce BCA Protein Assay Kit (Thermo Scientific). 10 µg of whole cell lysates or 2 µg of Nuclear Fractions were boiled in loading buffer (NuPAGE LDS Sample Buffer; Thermo Scientific) with reducing agents (NuPAGE Sample Reducing Agent; Thermo Scientific). Samples were resolved on 4-12% Bis-Tris Protein Gels (Thermo Scientific) prior to transfer onto nitrocellulose membranes (Thermo Scientific). After blocking in 5% non-fat skim milk for 1 hour, membranes were incubated in primary antibodies overnight on a shaker at 4 °C.

After 3x washes in TBST (Pierce TBS Tween 20 Buffer; Thermo Scientific), HRP-conjugated secondary antibodies (Goat anti-Rabbit or anti-Mouse, Thermo Scientific) were applied at a 1:2000 dilution in TBST for 1 hour on a shaker at room temperature. After 3x washes in TBST, signals were detected with either a Pierce ECL Western Blotting Substrate kit (Thermo Scientific) or SuperSignal West Dura Extended Duration Substrate kit (Thermo Scientific) and visualized with a ChemiDoc XRS+ imaging system equipped with Image Lab Software (Bio-Rad).

The following antibodies, diluted at the indicated concentrations in TBST supplemented with 3% Bovine Serum Albumin (Equitech-Bio), were used: ERα (1:1000), H3K9ac (1:9000), H3K27ac (1:500), H3K4me3 (1:1000), H3K9me3 (1:2000), H3K27me3 (1:500), H3 (1:3000), beta-Actin (1:3000) (Cell Signaling Technology), and H3K4ac (1:2000) (EMD Millipore).

### RNA isolation, reverse transcription, and real-time PCR

Total RNA was isolated from tissue culture plates according to the TRIzol Reagent (Invitrogen) protocol. 3 µg of total RNA was used in the reverse transcription reaction using the iScript cDNA synthesis kit (Bio-Rad). Quantitative PCR amplification was performed on the Prism 7900 Sequence Detection System (Applied Biosystems) using Taqman Gene Expression Assays (Applied Biosystems). Gene expression data were normalized to 18S rRNA.

### RNA Sequencing

Total RNA from two cell lines under two culture conditions (MDA-MB-231 +Ser, MDA-MB-231 -Ser, MCF7 +Ser, MCF7 -Ser) was extracted using the TRIzol Reagent (Invitrogen) protocol (n = 3 per group). The RNA-seq library was constructed and subjected to 150 base-pair paired-end sequencing on an Illumina sequencing platform (Novogene). RNA-seq analysis was performed using the kallisto and sleuth analytical pipeline^37,38^. In brief, a transcript index was generated with reference to Ensembl version 67 for hg19. Paired-end mRNA-seq reads were pseudo-aligned using kallisto (v0.42.4) with respect to this transcript index using 100 bootstraps (-b 100) to estimate the variance of estimated transcript abundances. Transcript-level estimates were aggregated to transcripts per million (TPM) estimates for each gene, with gene names assigned from Ensembl using biomaRt. Differential gene expression analysis was performed using the sleuth R package comparing the +Ser to -Ser condition of each cell line. Significant differentially expressed genes were identified using the Wald test function with cutoffs of q-value < 0.05 and fold change estimate b > abs(ln(2)).

### ChIP-Seq analysis

Chromatin immunoprecipitation followed by sequencing (ChIP-Seq) data in MCF7 cells were downloaded from the NCBI database (GSE130852). Peak calling was performed using MACS2 with peak detection cutoff based on p = 1 × 10^−5^. Peak annotation was performed in ChIPseeker in R, with enrichment calculated within a ± 3Kb region relative to each transcription start site. Individual tracks were visualized in igvtools.

### Metabolic tracing

5 × 10^5^ cells MCF7 cells were plated in 60 mm dishes overnight in DMEM/F12. The next day, the media was replaced with glucose/serine/glycine-free RPMI media (Teknova) supplemented with 10% dialyzed fetal bovine serum (Sigma) and 2.0 g/L ^13^C6-glucose (Sigma) to reconstitute - Ser media, and with 0.3 mM serine to reconstitute +Ser media, for 6 hours (n = 3 per group).

### Metabolite harvesting and liquid chromatography-mass spectrometry analysis

Cells were washed with cold PBS, lysed in 80% Ultra LC-MS acetonitrile (Thermo Scientific) on ice for 15 minutes, and centrifuged for 10 minutes at 20,000 x g at 4 °C. 200 µL of supernatants were subjected to mass spectrometry analysis. Liquid chromatography was performed using an Agilent 1290 Infinity LC system (Agilent, Santa Clara, US) coupled to a Q-TOF 6545 mass spectrometer (Agilent, Santa Clara, US). A hydrophilic interaction chromatography method with a ZIC-pHILIC column (150 × 2.1 mm, 5 µm; EMD Millipore) was used for compound separation at 35 °C with a flow rate of 0.3 mL/min. Mobile phase A consisted of 25 mM ammonium carbonate in water and mobile phase B was acetonitrile. The gradient elution was 0—1.5 min, 80% B; 1.5—7 min, 80% B → 50% B, 7—8.5 min, 50% B; 8.5—8.7 min, 50% B → 80% B; 8.7-13 min, 80% B. The overall runtime was 13 minutes, and the injection volume was 5 µL. The Agilent Q-TOF was operated in negative mode and the relevant parameters were as listed: ion spray voltage, 3500 V; nozzle voltage, 1000 V; fragmentor voltage, 125 V; drying gas flow, 11 L/min; capillary temperature, 325 °C; drying gas temperature, 350 °C; and nebulizer pressure, 40 psi. A full scan range was set at 50 to 1600 (m/z). The reference masses were 119.0363 and 980.0164. The acquisition rate was 2 spectra/s. Targeted analysis, isotopologues extraction (for the metabolic tracing study), and natural isotope abundance correction were performed by the Agilent Profinder B.10.00 Software (Agilent Technologies).

### Seahorse mitochondrial stress test

Oxygen consumption rates (OCR) were measured using the Seahorse XF Extracellular Flux Analyzer (Agilent Technologies, Santa Clara). MCF7 cells were plated at 1 × 10^4^ cells per well into the Seahorse XF96 plate in DMEM/F12 overnight. The next day, the media was replaced with +/- Ser media and pre-equilibrated for 2 hours at 37 °C. Mitochondrial stress tests were performed according to manufacturers’ protocol. After baseline OCR measurements, oligomycin A at 2.5 µM, trifluoromethoxy carbonylcyanide phenylhydrazone (FCCP) at 5 µM, and rotenone/antimycin A at 0.5/0.5 µM were sequentially injected into each well. All OCR measurements were normalized post-run with well-by-well cell counts on a hemocytometer.

### Quantification and statistical analysis

Statistical analyses were performed using GraphPad Prism 8.0. Groups of n = 3 were analyzed using the two-tailed students t test. Volcano plots were plotted using the EnhancedVolcano package in R. All error bars represent mean with standard deviation, and symbols denoting p-value significance are explained in the corresponding figure legends.

## KEY RESOURCES TABLE

| REAGENT or RESOURCE | SOURCE | IDENTIFIER |
| --- | --- | --- |
| Antibodies |  |  |
| Anti-ERα | Cell Signaling Technologies | 8644S |
| Anti-H3K4ac | EMD Millipore | 07-539 |
| Anti-H3K9ac | Cell Signaling Technologies | 9649S |
| Anti-H3K27ac | Cell Signaling Technologies | 8173S |
| Anti-H3K4me3 | Cell Signaling Technologies | 9727S |
| Anti-H3K9me3 | Cell Signaling Technologies | 13969S |
| Anti-H3K27me3 | Cell Signaling Technologies | 9733S |
| Anti-H3 | Cell Signaling Technologies | 3638S |
| Anti-beta-Actin | Cell Signaling Technologies | 3700S |
| Goat anti-Mouse IgG (H+L), HRP | Thermo Scientific | 62-652-0 |
| Goat anti-Rabbit IgG (H+L), HRP | Thermo Scientific | 65-6120 |
| Chemicals, Peptides, and Recombinant Proteins |  |  |
| Gibco-Penicillin-Streptomycin (10,000 U/mL) | Thermo Scientific | 15140122 |
| Trypan Blue Stain | Thermo Scientific | 15250061 |
| Fulvestrant | Cayman | 10011269 |
| Tamoxifen, white powder, Packaging=Amber Glass Bottle | MP Biomedicals | 0215673880 |
| RIPA Lysis and Extraction Buffer | Thermo Scientific | 89900 |
| Halt Protease and Phosphatase Inhibitor Single-Use Cocktail | Thermo Scientific | 78442 |
| NuPAGE LDS Sample Buffer (4X) | Thermo Scientific | NP0007 |
| NuPAGE Sample Reducing Agent (10X) | Thermo Scientific | NP0004 |
| TRIzol Reagent | Thermo Scientific | 15596026 |
| D-Glucose-<sup>13</sup>C<sub>6</sub> | Sigma | 389374-1G |
| Acetonitrile, Optima LC/MS Grade, Fisher Chemical | Thermo Scientific | A955-4 |
| Oligomycin A, ≥95% (HPLC) | Sigma | 75351-5MG |
| Carbonyl cyanide 4-(trifluoromethoxy)phenylhydrazone | Sigma | C2920-10MG |
| Rotenone | Sigma | R8875-1G |
| Antimycin A from *Streptomyces* sp. | Sigma | A8674-25MG |
| Critical Commercial Assays |  |  |
| Pierce BCA Protein Assay kit | Thermo Scientific | 23225 |
| Pierce ECL Western Blotting Substrate kit | Thermo Scientific | 32106 |
| SuperSignal West Dura Extended Duration Substrate kit | Thermo Scientific | PIA34075 |
| iScript cDNA Synthesis kit | Bio-Rad | 1708891 |
| Deposited Data |  |  |
| ChIP-seq (MCF7 cells) | GEO | GSE130852 |
| Experimental Models: Cell Lines |  |  |
| MCF7 | ATCC | HTB-22 |
| T47D | ATCC | HTB-133 |
| MDA-MB-231-Parental | MSKCC | RRID:CVCL_VR35 |
| MDA231-LM2-4175 | MSKCC | RRID:CVCL_5998 |
| MDA-BoM-1833 | MSKCC | RRID:CVCL_DP48 |
| Oligonucleotides |  |  |
| TaqMan probe human ESR1 | Thermo Scientific | Hs06598923_s1 |
| TaqMan probe human ESR2 | Thermo Scientific | Hs01100353_m1 |
| TaqMan probe human GPER1 | Thermo Scientific | Hs01922715_s1 |
| TaqMan probe human PGR | Thermo Scientific | Hs01556702_m1 |
| TaqMan probe human SLC1A4 | Thermo Scientific | Hs00983079_m1 |
| TaqMan probe human SLC1A5 | Thermo Scientific | Hs01056542_m1 |
| TaqMan probe human ASNS | Thermo Scientific | Hs04186194_m1 |
| TaqMan probe human PHGDH | Thermo Scientific | Hs00198333_m1 |
| TaqMan probe human PSAT1 | Thermo Scientific | Hs00795278_mH |
| TaqMan probe human PSPH | Thermo Scientific | Hs00190154_m1 |
| TaqMan probe human SHMT1 | Thermo Scientific | Hs00541045_m1 |
| TaqMan probe human SHMT2 | Thermo Scientific | Hs01059263_g1 |
| TaqMan probe human MTHFD2 | Thermo Scientific | Hs00759197_s1 |
| TaqMan probe human MTHFD1L | Thermo Scientific | Hs00914916_m1 |
| TaqMan probe human 18S | Thermo Scientific | Hs99999901_s1 |
| Software and Algorithms |  |  |
| Image Lab | Bio-Rad | N/A |
| GraphPad Prism 8.0 | GraphPad Software | N/A |
| ENRICHR | Ma'ayan Lab, Icahn School of Medicine at Mount Sinai, Center for Bioinformatics | N/A |
| GSEA_4.1.0 | Broad Institute | N/A |

---

**Supplementary Figure S1.**
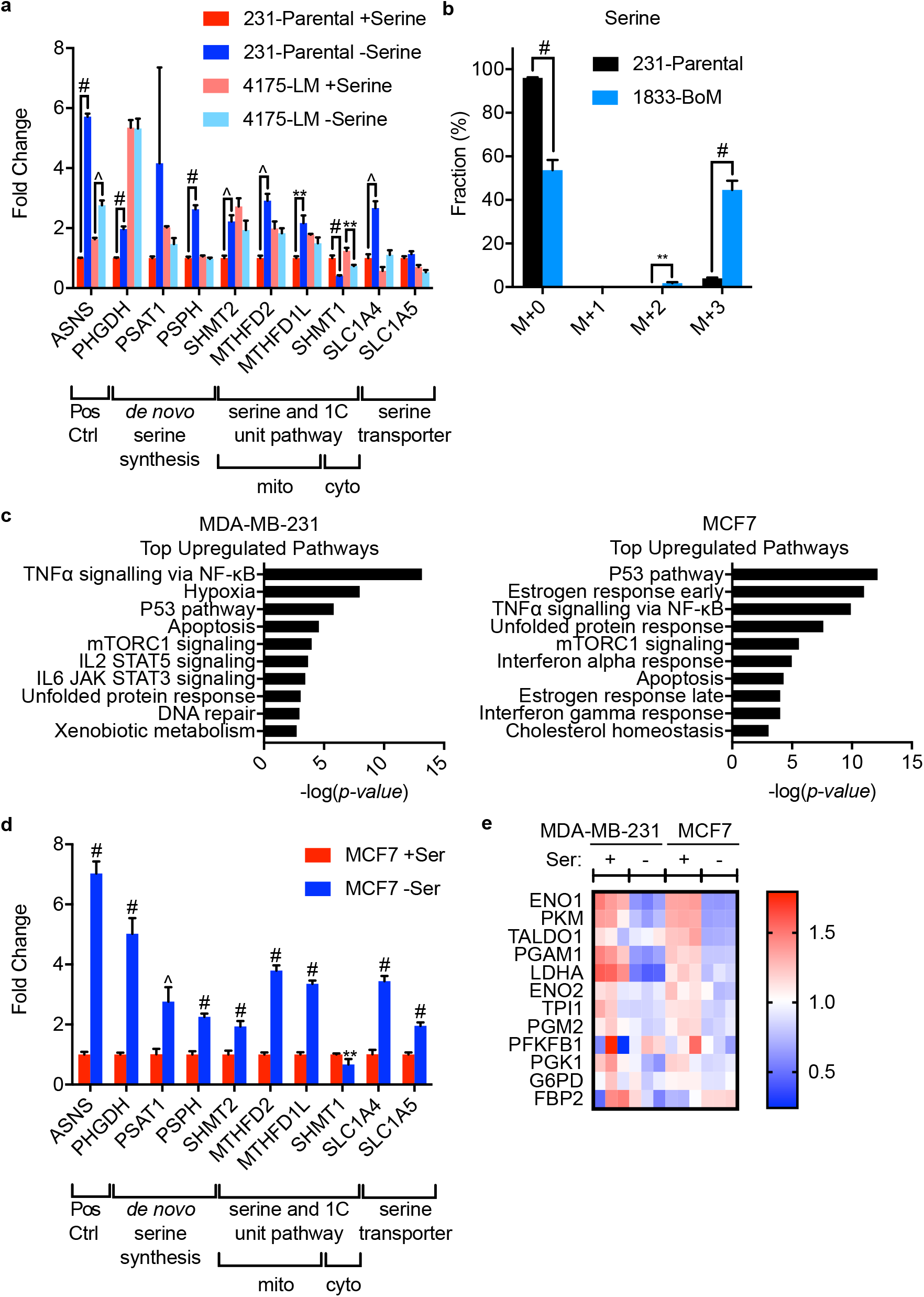
Transcriptional changes and metabolic reprogramming of breast cancer cells under serine starvation. **(a)** Quantitative RT-PCR for genes involved in serine metabolism (synthesis, catabolism, and import) from parental MDA-MB-231 (231-parental) and lung metastatic (4175-LM) cells. **(b)** Isotopomer distribution of serine from 231-parental or bone metastatic (1833-BoM) cells cultured in complete or serine-free, glucose-free media supplemented with U-^13^C-glucose for 6h. **(c)** Pathway analysis of top upregulated pathways via ENRICHR performed in control vs. 24h serine-starved MDA-MB-231 or MCF7 cells. **(d)** Quantitative RT-PCR for genes involved in serine metabolism (synthesis, catabolism, and import) from MCF7 cells. **(e)** Heatmap of the expression of genes encoding select glycolytic enzymes as measured by RNA-seq. Data are mean ± SD **(a, b, d)**, N = 3 per group. Statistics: two-tailed unpaired Student’s t test; **p < 0.01, ^p < 0.001, #p < 0.0001.

**Supplementary Figure S2.**
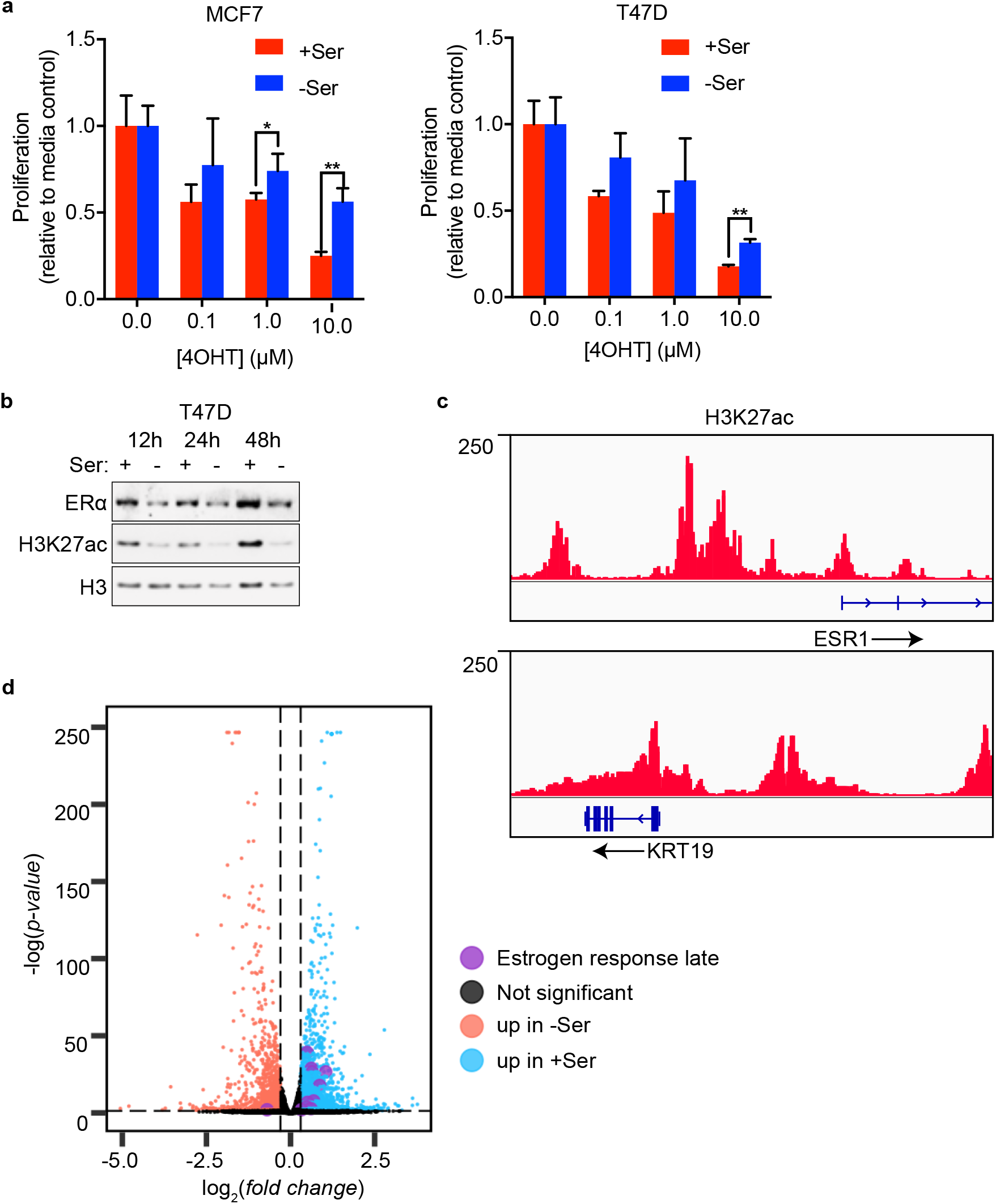
ER^+^ cell line phenotypes under serine starvation. **(a)** Four day proliferation assay on MCF7 and T47D cells in control (+Ser) or serine-free (-Ser) media, treated with tamoxifen (4OHT). **(b)** Immunoblot against estrogen receptor alpha (ERα) and H3K27ac, performed on nuclear fraction lysates isolated from T47D cells grown in +/- Ser media at 12h, 24h, and 48h. **(c)** MCF7 H3K27ac levels determined by ChIP-seq analysis at the *ESR1* and *KRT19* genes. **(d)** Volcano plot of DEGs obtained from RNA-seq in serine-starved MCF7 cells. Purple dots represent genes belonging to the estrogen response late pathway that also contain at least one H3K27ac peak. Data are mean ± SD **(a)**, N = 3 per group. Statistics: two-tailed unpaired Student’s t test; *p < 0.05, **p < 0.01.

**Supplementary Figure S3.**
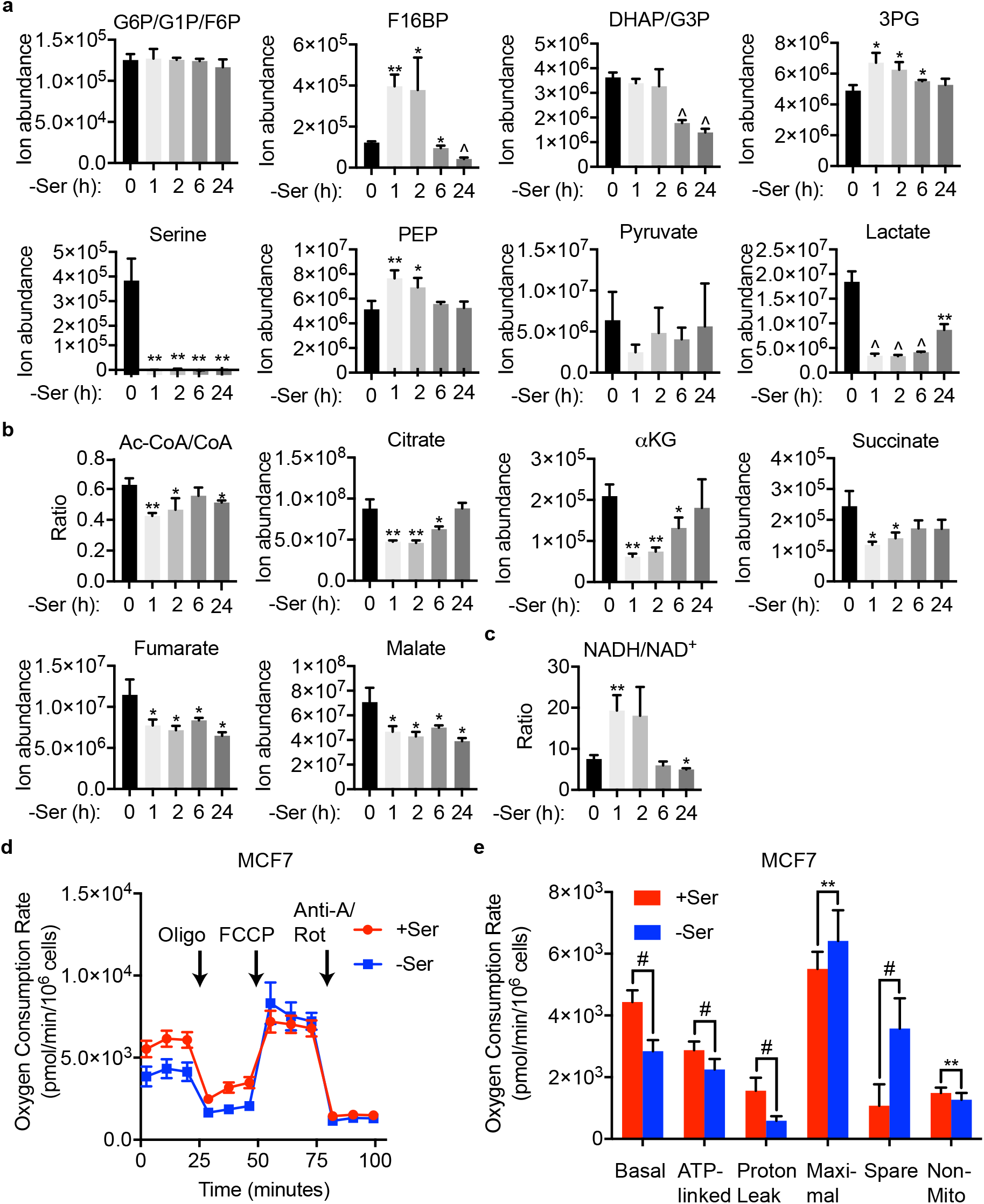
Kinetic analysis of metabolic changes in ER^+^ breast cancer cells under serine starvation. **(a-c)** LC-MS was used to determine the levels of intermediates in glycolysis **(a)**, the TCA cycle **(b)**, and the NADH/NAD^+^ ratio **(c)** at various time points of serine starvation in T47D cells. Abbreviations: G6P, glucose 6-phosphate; G1P, glucose 1-phosphate; F6P, fructose 6-phoshpate; F16BP, fructose 1,6-bisphosphate; DHAP, dihydroxyacetone phosphate; G3P, glycerol 3-phoshpate; 3PG, 3-phosphoglycerate; PEP, phosphoenolpyruvate; Ac-CoA, acetyl-co enzyme A; CoA, co-enzyme A; αKG, alpha ketoglutarate; NAD^+^, nicotinamide adenine dinucleotide. **(d)** Seahorse assay for the oxygen consumption rate (OCR) of MCF7 cells cultured in +/- Ser in real time under basal conditions and in response to mitochondrial inhibitors. Abbreviations: Oligo, oligomycin; FCCP, trifluoromethoxy carbonylcyanide phenylhydrazone; Anti-A, antimycin A; Rot, rotenone. **(e)** Calculated values for respiratory parameters collected from **(d)**. Abbreviations: Basal, basal respiration; ATP-linked, ATP-linked respiration; Maximal, maximal respiratory capacity; Spare, spare respiratory capacity; Non-mito, non-mitochondrial respiration. Data are mean ± SD **(a-c, e)**, N = 3 per group **(a-c)** and N = 5 per group **(d, e)**. Statistics: two-tailed unpaired Student’s t test; *p < 0.05, **p < 0.01, ^p < 0.001, #p < 0.0001.

## REFERENCES

1 Osborne, C. K. & Schiff, R. Mechanisms of Endocrine Resistance in Breast Cancer. Annu Rev Med 62, 233–247, doi:10.1146/annurev-med-070909-182917 (2011).

2 Chang, M. Tamoxifen resistance in breast cancer. Biomol Ther (Seoul) 20, 256–267, doi:10.4062/biomolther.2012.20.3.256 (2012).

3 Lim, E. et al. Aberrant luminal progenitors as the candidate target population for basal tumor development in BRCA1 mutation carriers. Nat Med 15, 907–913, doi:10.1038/nm.2000 (2009).

4 Fan, M. et al. Triggering a switch from basal-to luminal-like breast cancer subtype by the small-molecule diptoindonesin G via induction of GABARAPL1. Cell Death Dis 11, 635, doi:10.1038/s41419-020-02878-z (2020).

5 Pavlova, N. N. & Thompson, C. B. The Emerging Hallmarks of Cancer Metabolism. Cell Metab 23, 27–47, doi:10.1016/j.cmet.2015.12.006 (2016).

6 Lukey, M. J. et al. Liver-Type Glutaminase GLS2 Is a Druggable Metabolic Node in Luminal-Subtype Breast Cancer. Cell Rep 29, 76–88 e77, doi:10.1016/j.celrep.2019.08.076 (2019).

7 Li, A. M. et al. Metabolic Profiling Reveals a Dependency of Human Metastatic Breast Cancer on Mitochondrial Serine and One-Carbon Unit Metabolism. Mol Cancer Res, doi:10.1158/1541-7786.MCR-19-0606 (2020).

8 Sun, X. et al. Metabolic Reprogramming in Triple-Negative Breast Cancer. Front Oncol 10, 428, doi:10.3389/fonc.2020.00428 (2020).

9 Mahendralingam, M. J. et al. Mammary epithelial cells have lineage-rooted metabolic identities. Nat Metab 3, 665–681, doi:10.1038/s42255-021-00388-6 (2021).

10 Pan, M. et al. Regional glutamine deficiency in tumours promotes dedifferentiation through inhibition of histone demethylation. Nat Cell Biol 18, 1090–1101, doi:10.1038/ncb3410 (2016).

11 Kamphorst, J. J. et al. Human pancreatic cancer tumors are nutrient poor and tumor cells actively scavenge extracellular protein. Cancer Res 75, 544–553, doi:10.1158/0008-5472.CAN-14-2211 (2015).

12 Sullivan, M. R. et al. Increased Serine Synthesis Provides an Advantage for Tumors Arising in Tissues Where Serine Levels Are Limiting. Cell Metab 29, 1410–1421 e1414, doi:10.1016/j.cmet.2019.02.015 (2019).

13 Yang, M. & Vousden, K. H. Serine and one-carbon metabolism in cancer. Nat Rev Cancer 16, 650–662, doi:10.1038/nrc.2016.81 (2016).

14 Maddocks, O. D. et al. Serine starvation induces stress and p53-dependent metabolic remodelling in cancer cells. Nature 493, 542–546, doi:10.1038/nature11743 (2013).

15 Maddocks, O. D. K. et al. Modulating the therapeutic response of tumours to dietary serine and glycine starvation. Nature 544, 372–376, doi:10.1038/nature22056 (2017).

16 Ye, J. et al. Pyruvate kinase M2 promotes de novo serine synthesis to sustain mTORC1 activity and cell proliferation. Proc Natl Acad Sci U S A 109, 6904–6909, doi:10.1073/pnas.1204176109 (2012).

17 Possemato, R. et al. Functional genomics reveal that the serine synthesis pathway is essential in breast cancer. Nature 476, 346–U119, doi:10.1038/nature10350 (2011).

18 Choi, B. H., Conger, K. O., Selfors, L. M. & Coloff, J. L. Lineage-specific silencing of PSAT1 induces serine auxotrophy and sensitivity to dietary serine starvation in luminal breast tumors. bioRxiv, doi:https://doi.org/10.1101/2020.06.19.161844 (2020).

19 Kang, Y. et al. A multigenic program mediating breast cancer metastasis to bone. Cancer Cell 3, 537–549, doi:10.1016/s1535-6108(03)00132-6 (2003).

20 Minn, A. J. et al. Genes that mediate breast cancer metastasis to lung. Nature 436, 518–524, doi:10.1038/nature03799 (2005).

21 Bos, P. D. et al. Genes that mediate breast cancer metastasis to the brain. Nature 459, 1005–1009, doi:10.1038/nature08021 (2009).

22 Booms, A., Coetzee, G. A. & Pierce, S. E. MCF-7 as a Model for Functional Analysis of Breast Cancer Risk Variants. Cancer Epidemiol Biomarkers Prev 28, 1735–1745, doi:10.1158/1055-9965.EPI-19-0066 (2019).

23 Lu, C. & Thompson, C. B. Metabolic regulation of epigenetics. Cell Metab 16, 9–17, doi:10.1016/j.cmet.2012.06.001 (2012).

24 Reid, M. A., Dai, Z. & Locasale, J. W. The impact of cellular metabolism on chromatin dynamics and epigenetics. Nat Cell Biol 19, 1298–1306, doi:10.1038/ncb3629 (2017).

25 Schvartzman, J. M., Thompson, C. B. & Finley, L. W. S. Metabolic regulation of chromatin modifications and gene expression. J Cell Biol 217, 2247–2259, doi:10.1083/jcb.201803061 (2018).

26 Wellen, K. E. et al. ATP-citrate lyase links cellular metabolism to histone acetylation. Science 324, 1076–1080, doi:10.1126/science.1164097 (2009).

27 Diehl, F. F., Lewis, C. A., Fiske, B. P. & Vander Heiden, M. G. Cellular redox state constrains serine synthesis and nucleotide production to impact cell proliferation. Nature Metabolism 1, 861–867, doi:10.1038/s42255-019-0108-x (2019).

28 Peng, M. et al. Aerobic glycolysis promotes T helper 1 cell differentiation through an epigenetic mechanism. Science 354, 481–484, doi:10.1126/science.aaf6284 (2016).

29 Gao, X. et al. Acetate functions as an epigenetic metabolite to promote lipid synthesis under hypoxia. Nat Commun 7, 11960, doi:10.1038/ncomms11960 (2016).

30 Li, Y. et al. Acetate supplementation restores chromatin accessibility and promotes tumor cell differentiation under hypoxia. Cell Death Dis 11, 102, doi:10.1038/s41419-020-2303-9 (2020).

31 Ngo, B. et al. Limited Environmental Serine and Glycine Confer Brain Metastasis Sensitivity to PHGDH Inhibition. Cancer Discov 10, 1352–1373, doi:10.1158/2159-8290.CD-19-1228 (2020).

32 Rinaldi, G. et al. In Vivo Evidence for Serine Biosynthesis-Defined Sensitivity of Lung Metastasis, but Not of Primary Breast Tumors, to mTORC1 Inhibition. Molecular Cell 81, 386-+, doi:10.1016/j.molcel.2020.11.027 (2021).

33 Chaneton, B. et al. Serine is a natural ligand and allosteric activator of pyruvate kinase M2. Nature 491, 458–462, doi:10.1038/nature11540 (2012).

34 Kurebayashi, J., Otsuki, T., Moriya, T. & Sonoo, H. Hypoxia reduces hormone responsiveness of human breast cancer cells. Jpn J Cancer Res 92, 1093–1101, doi:DOI 10.1111/j.1349-7006.2001.tb01064.x (2001).

35 Murphy, J. P. et al. The NAD(+) Salvage Pathway Supports PHGDH-Driven Serine Biosynthesis. Cell Rep 24, 2381–2391 e2385, doi:10.1016/j.celrep.2018.07.086 (2018).

36 Yang, X. W. et al. Synergistic activation of functional estrogen receptor (ER)-alpha by DNA methyltransferase and histone deacetylase inhibition in human ER-alpha-negative breast cancer cells. Cancer Research 61, 7025–7029 (2001).

37 Bray, N. L., Pimentel, H., Melsted, P. & Pachter, L. Near-optimal probabilistic RNA-seq quantification. Nat Biotechnol 34, 525–527, doi:10.1038/nbt.3519 (2016).

38 Pimentel, H., Bray, N. L., Puente, S., Melsted, P. & Pachter, L. Differential analysis of RNA-seq incorporating quantification uncertainty. Nat Methods 14, 687-+, doi:10.1038/nmeth.4324 (2017).

